# Enzyme Catalysis Enhances Lateral Lipid Motility and Directional Particle Transport on Membranes

**DOI:** 10.1101/2021.09.06.459117

**Authors:** Ambika Somasundar, Niladri Sekhar Mandal, Ayusman Sen

## Abstract

The dynamic interplay between the composition of lipid membranes and the behavior of membrane-bound enzymes is critical to the understanding of cellular function and viability, and the design of membrane-based biosensing platforms. While there is a significant body of knowledge on how lipid composition and dynamics affect membrane-bound enzymes, little is known about how enzyme catalysis influences the motility and lateral transport in lipid membranes. Using enzymes-attached lipids in supported bilayers (SLB), we show catalysis-induced enhanced lateral diffusion of lipids in the bilayer. Enhancing the membrane viscosity by increasing the cholesterol content in the bilayer suppresses the overall diffusion but not the relative diffusion enhancement of the enzyme-attached lipids. We also provide direct evidence of catalysis-induced membrane fluctuations leading to the enhanced diffusion of passive tracers resting on the SLB. Additionally, by using active enzyme patches, we demonstrate the directional transport of tracers on SLBs. These are first steps in understanding diffusion and transport in lipid membranes due to active, out-of-equilibrium processes that are the hallmark of living systems. In general, our study demonstrates how active enzymes can be used to control diffusion and transport in confined 2-D environments.

---

Lipid membranes are critical to the viability of living systems,^1^ and the dynamic motility and transient spatial reorganization of membrane components is important for many essential cellular processes.^2^ While lipids provided the basic structural framework, the membrane-bound enzymes carry out the vast array of functions from molecular transport to regulation of signaling pathways. Membrane-bound enzymes are also important targets for pathogens and drugs,^3^ and are the basis for biosensing platforms.^4^ Therefore, it is essential to understand the dynamic interplay between lipid motility and enzyme catalysis in membranes. While there is a significant body of knowledge on how lipid composition and dynamics affect membrane-bound enzymes,^5,6^ little is known, regarding the effect of enzyme catalysis on the motility and lateral transport in lipid bilayers in cells.^7^

Due to the high complexity and heterogeneity of cell membranes, it is difficult to identify the specific mechanisms for lateral mobility and spatial reorganization in live cell membranes. Synthetic lipid membranes prepared with fewer membrane components can serve as useful models.^8,9^ Some of the commonly studied model systems include lipid vesicles, solid lipid particles, lipid monolayers, and supported lipid bilayers. Lipid membrane fluctuations and spontaneous pattern formation have been reported on ternary lipid systems.^10,11^ These studies mainly rely on lipids that spontaneously go into the most thermodynamically stable configuration. However, the ability to induce dynamic lipid fluctuations based on active or out-of-equilibrium processes such as catalysis is relatively unexplored. In this work, we probe the lateral mobility of lipids and membrane fluctuations by examining the diffusion coefficient of lipids tethered to active enzymes (**Figure 1**). We also study how the lateral diffusion is affected by the addition of passive membrane components such as cholesterol within the membrane or a polymer mimicking the extra cellular matrix above the membrane. We then examine how the diffusive behavior of inert tracer particles resting on lipid membrane changes due to catalysis-induced active motion of the lipid membranes (**Figure 1**). We also test the ability to direct passive particle transport on SLBs with asymmetric enzyme coverage by creating an enzyme-rich patch on the SLB (**Figure 1**).

**Figure 1.**
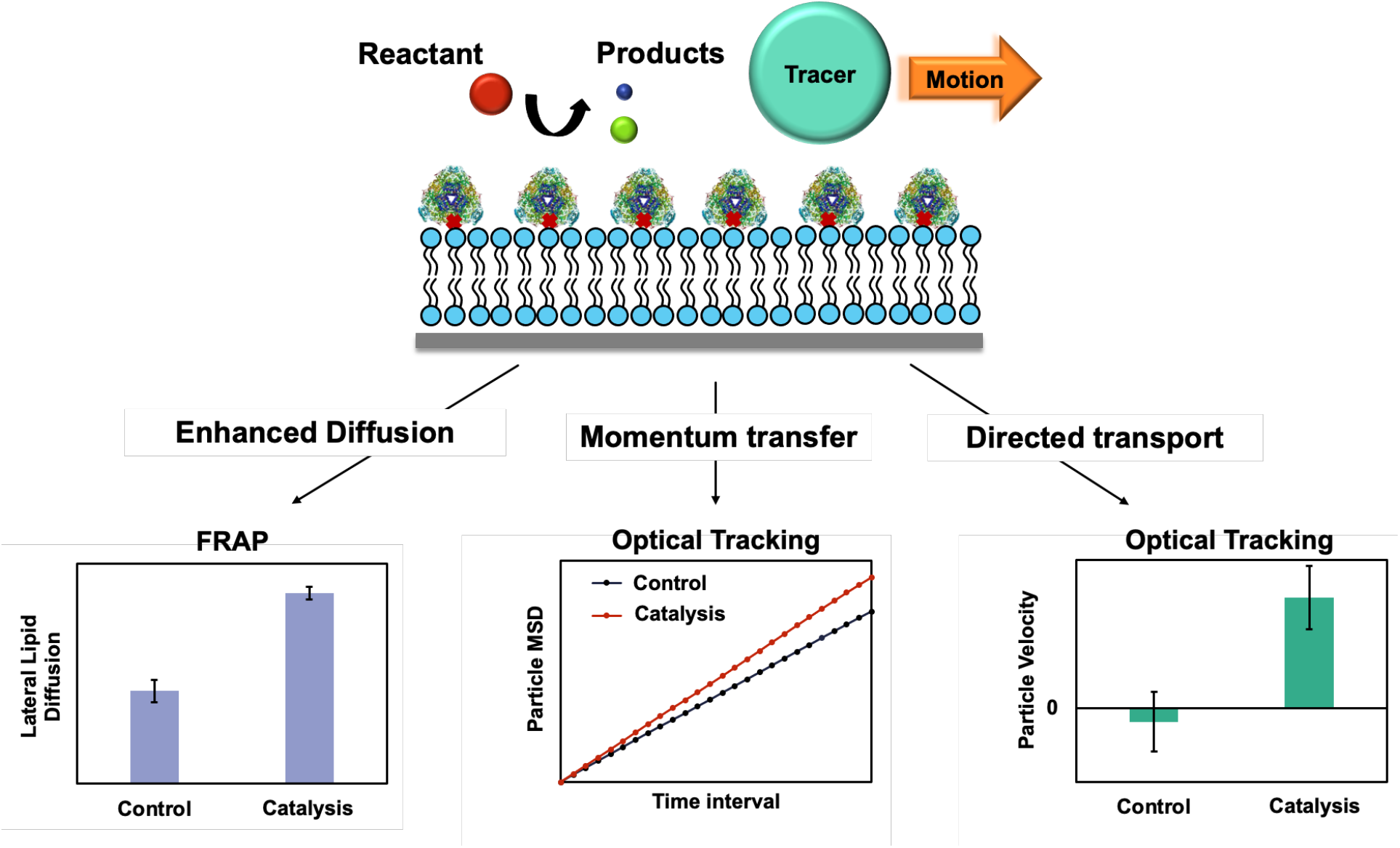
Schematic overview of enzyme-attached supported lipid bilayer (SLB). We observed enhanced diffusion of enzyme-attached lipids, catalytic momentum transfer to passive tracers and directional transport of passive tracers

It is known that the diffusive movement of enzymes in solution increases significantly during substrate turnover through the conversion of chemical energy into mechanical motion.^12–16^ This phenomenon, commonly termed as enhanced diffusion, has been used to power nano and microvehicles for theranostic applications. Attaching enzymes onto the surface of cargo particles such as polystyrene beads,^17^ polymersomes,^18,19^ or liposomes,^20,21^ enable them to be propelled directionally in gradients of their reactants. Here we show that the attachment of urease or hexokinase to confined surfaces such as a supported lipid bilayer (SLB) results in enhanced lateral motility of the enzyme-attached lipid upon catalysis. We also show that attenuating the fluidity of the membrane, through incorporation of cholesterol into the SLB does not change the relative increase in diffusion. This result is particularly significant in demonstrating that enhanced diffusion remains relevant in highly crowded or viscous cellular environment. We further show that spatially separating the enzyme from the lipid bilayer suppresses the enhanced lipid motility. This suggests that not only is active catalysis important for the enhanced mobility and fluctuations of membrane components, but also the way in which the components are associated in the membrane becomes important. Furthermore, the “momentum” arising from the enhanced diffusion of active enzyme-attached supported lipid bilayers can be transferred to passive tracers resting on the surface resulting in their increase in diffusive motion. This demonstrates the importance of active force fluctuations in membranes in enabling the motion of other cellular components. Controlling the motility of membrane components through out-of-equilibrium active process helps to understand the dynamic role they play in regulating key functions of the cell.

## Results and Discussion

### Increased diffusion of urease-attached lipids

We report the enhanced diffusion of enzymes tethered to SLBs via the Fluorescence Recovery After Photobleaching (FRAP) technique. To attach enzymes onto the SLB, we biotinylated the enzymes and tethered them to the biotinylated lipids via fluorescently tagged streptavidin (cy5) (**Figure 2**). Once the SLB is prepared, we bleach one region of the SLB and observe the rate of recovery of fluorescently tagged lipid. We then calculate the diffusion coefficient of the lipids by fitting the recovery curve with a simulated exponential recovery curve. Further experimental and measurement details are given in the Supplemental Information (SI).

**Figure 2.**
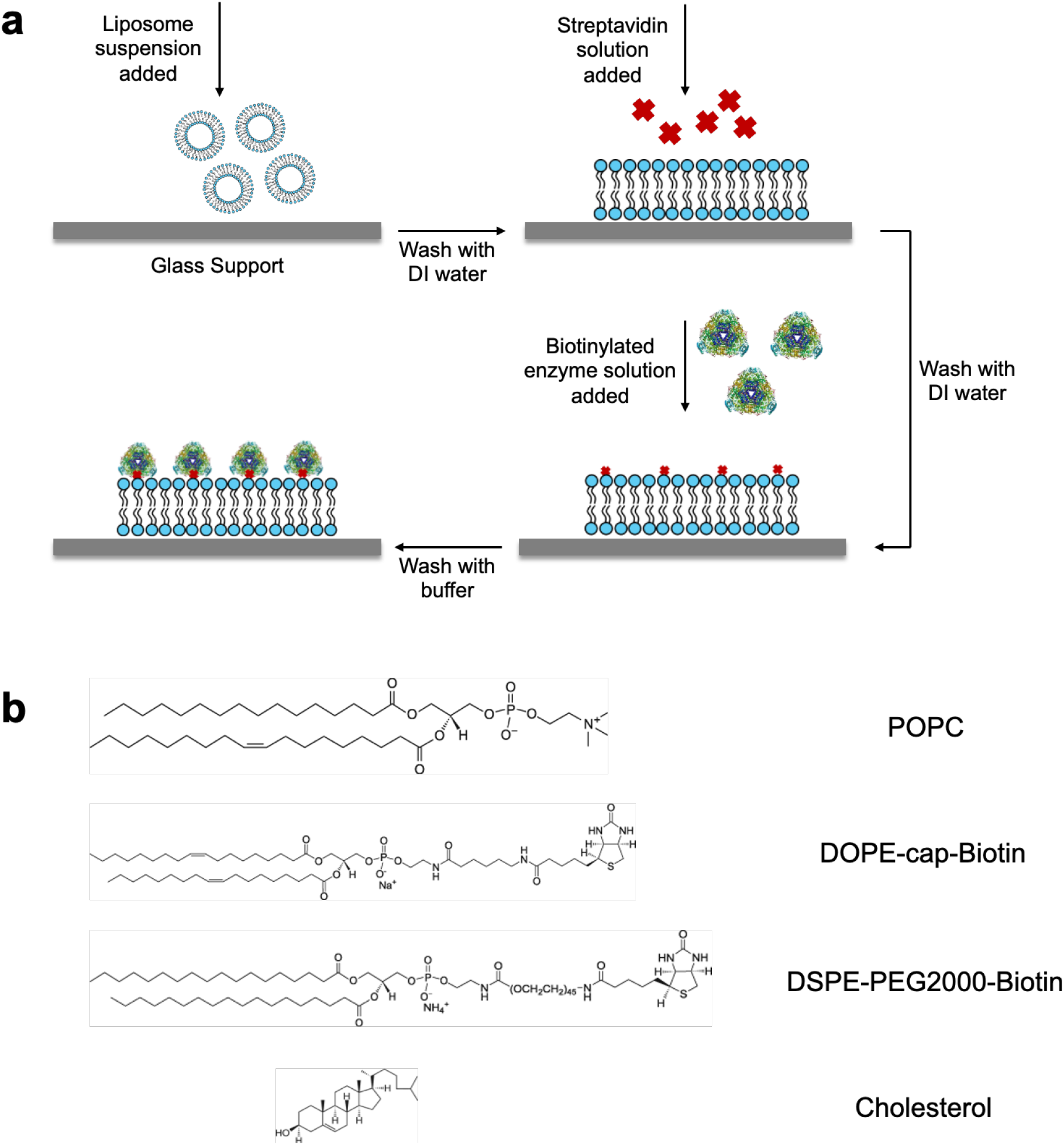
(a) Schematic and experimental procedure for enzyme-attached supported lipid bilayer (SLB) formation. (b) Structures of membrane components used for the preparation of SLBs.

The first enzyme we tested in this study is urease. Urease is a relatively fast enzyme that converts urea to ammonium and bicarbonate ions. It is also one of the enzymes whose enhanced diffusivity during catalysis has been verified by a variety of techniques, such as Fluorescence Correlation Spectroscopy (FCS)^12^, Dynamic Light Scattering (DLS)^17^, particle tracking^22^ and Total Internal Reflection Fluorescence Microscopy (TIRFM).^23^

We measured the mobility of the urease-attached lipid in the absence (control) and presence (experiment) of 200 mM urea. For the control case, we observe *D*_*buffer*_ = 0.084 µm^2^/s. This is a reasonable value, based on the expected two-dimensional viscosity at the membrane surface and the frictional coefficient that arises from the attachment of the lower leaflet of the SLB to the glass (see SI for details). We observed an increase in diffusion of urease upon catalysis to *D*_*urea*_ = 0.173 µm^2^/s (**Figure 3a)**. This corresponds to a 106% enhancement in diffusion of urease-attached lipid during catalysis. Interestingly, such a high enhancement in diffusion of urease has been observed only in surface confined experimental set up^23,24^ and not in freely diffusing enzyme solutions.^12,15^ In order to ensure that the enhancement of diffusion is indeed due to the urease reaction, we performed control experiments without the presence of urease to show that the addition of urea does not change the diffusion coefficient of the bilayer without catalysis (**Figure S1**).

**Figure 3.**
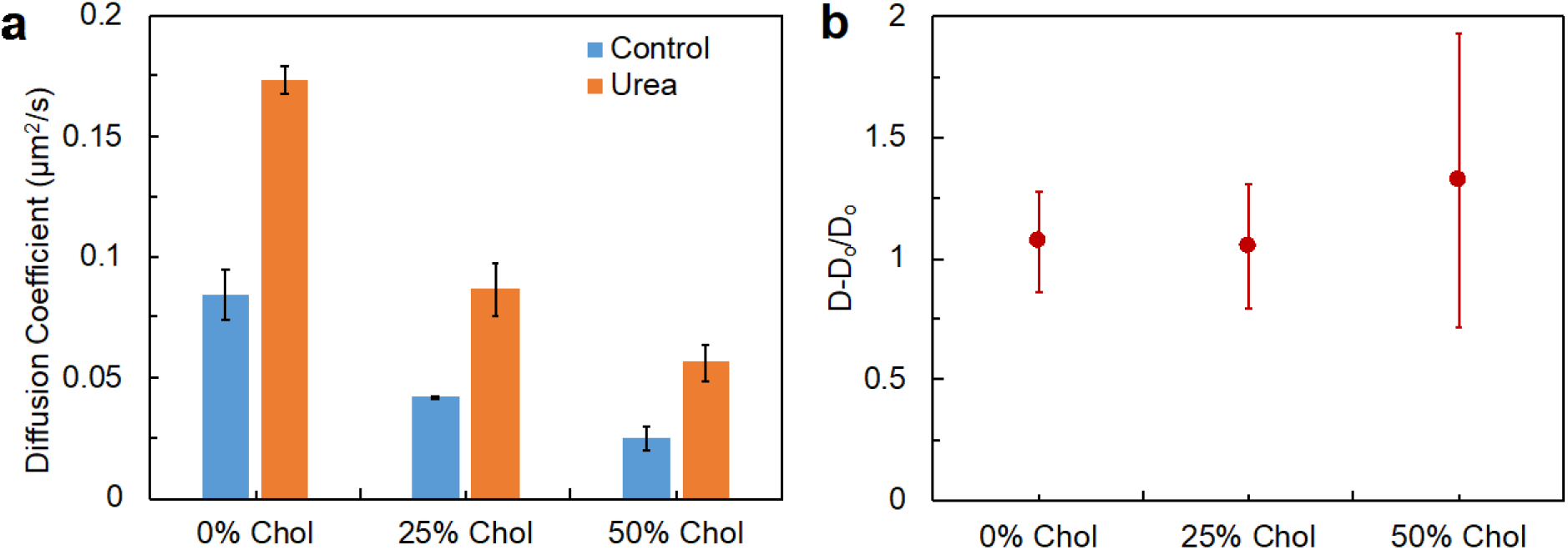
(a) Diffusion of urease-attached lipids with and without catalysis at 0, 25 and 50% cholesterol in SLB. Urease is attached to 1% Biotin-cap-DOPE in the SLB. (b) % enhancement in diffusion at 0, 25 and 50% cholesterol in SLB. The buffer used is 100 mM PBS at pH 7.2.

Next, we sought to increase the “viscosity” or rigidity of the SLB internally by adding cholesterol to the SLB itself and without altering the medium above it. We studied the enhanced diffusion of urease enzymes at cholesterol concentrations of 0, 25, and 50 mol% in the bilayer. We observed that while adding cholesterol to the bilayer decreases the baseline diffusivity of the urease-attached lipid (**Figure 3a**), the relative enhancement in diffusion still remains the same. The relative enhancement in diffusion for SLB with 25 and 50 mol% cholesterol are 105% and 134%, respectively (**Figure 3b**). While the relative enhancement in diffusion remains the same, the change in diffusion coefficient (ΔD) between the control and experiment decreases as the cholesterol content increases (**Figure S2a**). This is an important observation that will help to explain our observations in the momentum transfer experiments.

### Increased diffusion of hexokinase-attached lipids

To test the generality of enhanced diffusion of active lipid bilayers, we studied another enzyme, hexokinase. Hexokinase is a glycolytic enzyme that phosphorylates glucose to glucose-6-phosphate in the presence of ATP and MgCl_2_. In this case we observe that the relative enhancement of enzymes is ∼26%. For the control case, we observe *D*_*buffer*_ = 0.106 µm^2^/s and for the experiment, we observe *D*_*full catalysis*_ = 0.134 µm^2^/s (**Figure 4a**). For full catalysis, we use 20 mM D-glucose, 20 mM ATP and 40 mM MgCl_2_. We also performed control experiments without the presence of hexokinase to show that the reactants do not change the diffusion coefficient of the bilayer without catalysis (**Figure S3**).

**Figure 4.**
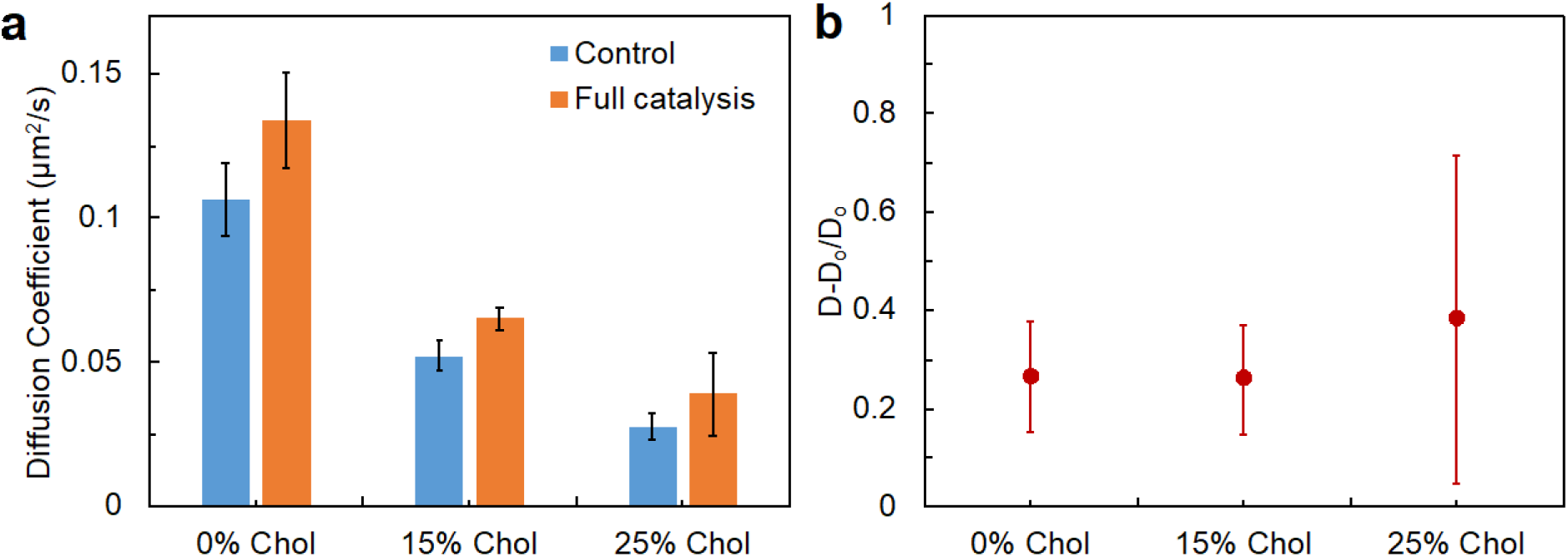
(a) Diffusion of hexokinase-attached lipids with and without catalysis at 0, 15 and 25% cholesterol in SLB. Hexokinase is attached to 1% Biotin-cap-DOPE in the SLB (b) % enhancement of diffusion at 0, 15 and 25% cholesterol in SLB. Full catalysis corresponds to 20 mM D-glucose, 20 mM ATP and 40 mM MgCl_2_. The buffer used is 50 mM HEPES at pH 7.4.

As with urease, as we increased the viscosity of the membrane by adding cholesterol into the SLB, we observed an attenuation of baseline diffusivity of the enzyme but not the relative enhancement in diffusivity upon catalysis (**Figure 4b**). In the case of hexokinase, we observed a steeper decrease in baseline diffusivity upon incorporation of cholesterol to the bilayer compared to that for urease. In the case of urease, the baseline diffusivity decreased from 0.08 µm^2^/s to 0.04 µm^2^/s when increasing cholesterol from 0 to 25 mol%. However, for hexokinase the baseline diffusivity goes from 0.1 µm^2^/s to 0.02 µm^2^/s when increasing the cholesterol concentration from 0 to 25 mol%. This suggests some interaction of hexokinase with cholesterol that promotes tighter binding of the enzyme to the bilayer, consistent with what has been reported previously.^25^ As with urease, we observed a decrease in the change in diffusion coefficient (ΔD) with increase in cholesterol concentration (**Figure S2b**).

### Long pegylated linkers suppress the diffusion enhancement by active enzymes

In cell membranes, apart from cholesterol, there are glycoproteins and glycolipids that rises tens of nanometers above the bilayer.^26^ These structures, collectively termed as glycocalyx, protect the cell from chemical injury, infection and adhesion among others.^27^ It has been proposed that a PEG coating serves as a simple mimic of the cell glycocalyx on supported phospholipid membranes.^28^ While we have demonstrated enhanced lipid diffusion with active enzymes close to the SLB, we now ask the question if the enhanced diffusion will also be observed if the active enzyme is further away from the SLB. To address this, we used a different biotin linker to attached enzymes to the SLB. Instead of Biotin-cap-DOPE, we used DSPE-PEG2000-Biotin (**Figure 5a**). With this longer linker, we observed no enhancement in the diffusion of the active urease attached lipid (**Figure 5b**). We hypothesize that the enzyme, being far away from the SLB, behaves like a free enzyme in solution, thereby preventing us from probing the diffusive behavior through the confocal FRAP technique.

**Figure 5.**
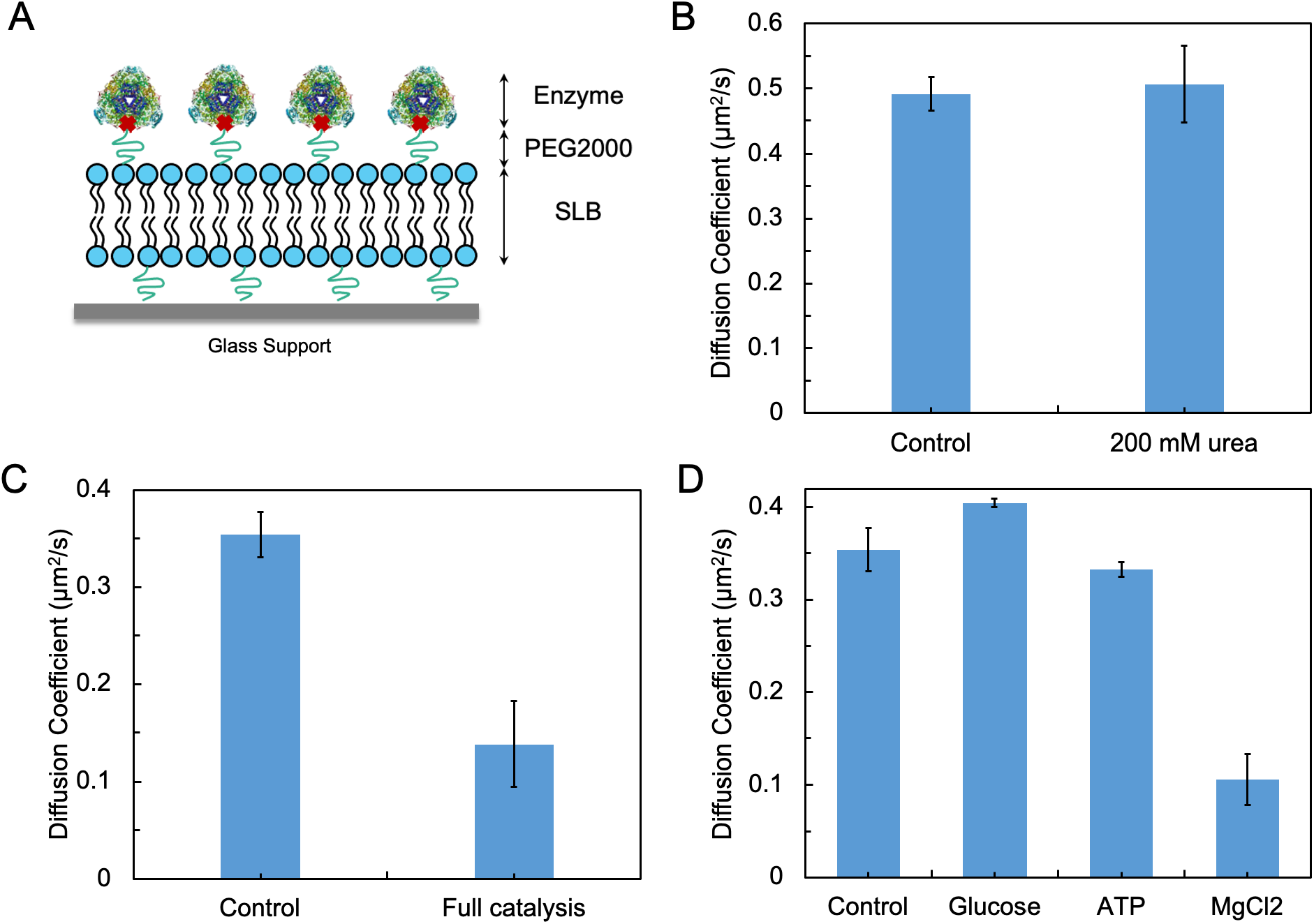
(a) Schematic representation of enzyme-attached SLB using a pegylated biotin linker. (b) Diffusion of urease-attached SLB with and without catalysis. The buffer used was 100 mM PBS at pH 7.2. (c) Diffusion of hexokinase-attached SLB with and without full catalysis. Full catalysis corresponds to 20 mM D-glucose, 20 mM ATP and 40 mM MgCl_2_. The buffer used is 50 mM HEPES at pH 7.4. (d) Diffusion of hexokinase-attached SLB in the presence of individual reactants required for catalysis. It is clear that MgCl_2_ is responsible for the freezing up of the bilayer. 1 mol% pegylated biotin linker was used in both the urease and hexokinase experiments.

A second interesting observation was that the baseline diffusion coefficient of the enzyme-attached lipid was five times higher for DSPE-PEG2000-Biotin, as compared to Biotin-cap-DOPE. In the absence of added cholesterol, the diffusion coefficients observed with DSPE-PEG2000-Biotin and Biotin-cap-DOPE were 0.49 µm^2^/s and 0.08 µm^2^/s, respectively. The increase in diffusion coefficient can be attributed to the long PEG2000 linker that increases the hydrophilic spacing between the glass support and the lower leaflet of the bilayer thus allowing the bilayer to diffuse more freely.^26,29^

With hexokinase attached to the SLB through the DSPE-PEG2000-Biotin linker, we observed that the diffusion coefficient was quenched upon full catalysis (**Figure 5c**). In order to understand the observed suppression of the lipid mobility, we performed control experiments with the individual reactants. When the hexokinase-attached SLB was exposed to either 20 mM D-glucose or 20 mM ATP, we did not see any dampening of diffusion coefficient. However, the addition of 40 mM MgCl_2_ (a co-factor required for catalytic activity) resulted in a sharp attenuation of the lipid diffusivity (**Figure 5d**). In contrast, with Biotin-cap-DOPE as the linker, the addition of MgCl_2_ did not result in diminution of lipid diffusivity. Thus, it is clear that the interaction of the Mg^2+^ ion with the PEG2000 chain is the reason for the “freezing” of the bilayer (See **Figure S4** for image of “frozen” SLB). We also tested the diffusion of just POPC SLB in the presence of MgCl_2_ and found that it does not have any effect on its diffusion (**Figure S5**).

### Enzyme-attached lipids transfer “momentum” to passive tracers on uniform enzyme-coated SLB surface

To further underscore the significance of catalysis-induced increased diffusion of lipid membranes, we examined the ability of the active bilayer to transfer “momentum” to passive tracers present on its surface. Previous studies have shown that enzymes in free solution are able to transfer “momentum” to surrounding tracer particles in solution.^30^ We determine here if this is also possible through catalysis-induced membrane fluctuations. For these experiments, we used urease-attached SLBs. The enzyme attached bilayers were prepared inside a hybridization chamber adhered to a cleaned glass coverslip (**Figure 6a**). BSA-coated 1 µm polystyrene tracer particles that were either mixed with buffer or the reactant urea were pipetted into the hybridization chamber. The chamber was then sealed from the top using a glass coverslip to prevent air bubbles and unwanted disturbances that can cause convective flows in the sample. The movement of the tracer particles on the plane closest to the bilayer were then captured using an inverted optical microscope. We ensured that the videos were taken right at the surface of the bilayer by observing that there were some particles adhered to the surface (and showing no Brownian motion) in the frame. This told us that we were in the plane closest to the bilayer. A minimum of 10 particles are tracked for each experiment and their trajectories were used to calculate MSD curves from MATLAB. Care was taken to ensure that only particles in videos that did not display convective flows were tracked. Additionally, the MATLAB code used accounted for any convective flows that could be present but were not be detected from the particle tracking videos in the sample.

**Figure 6.**
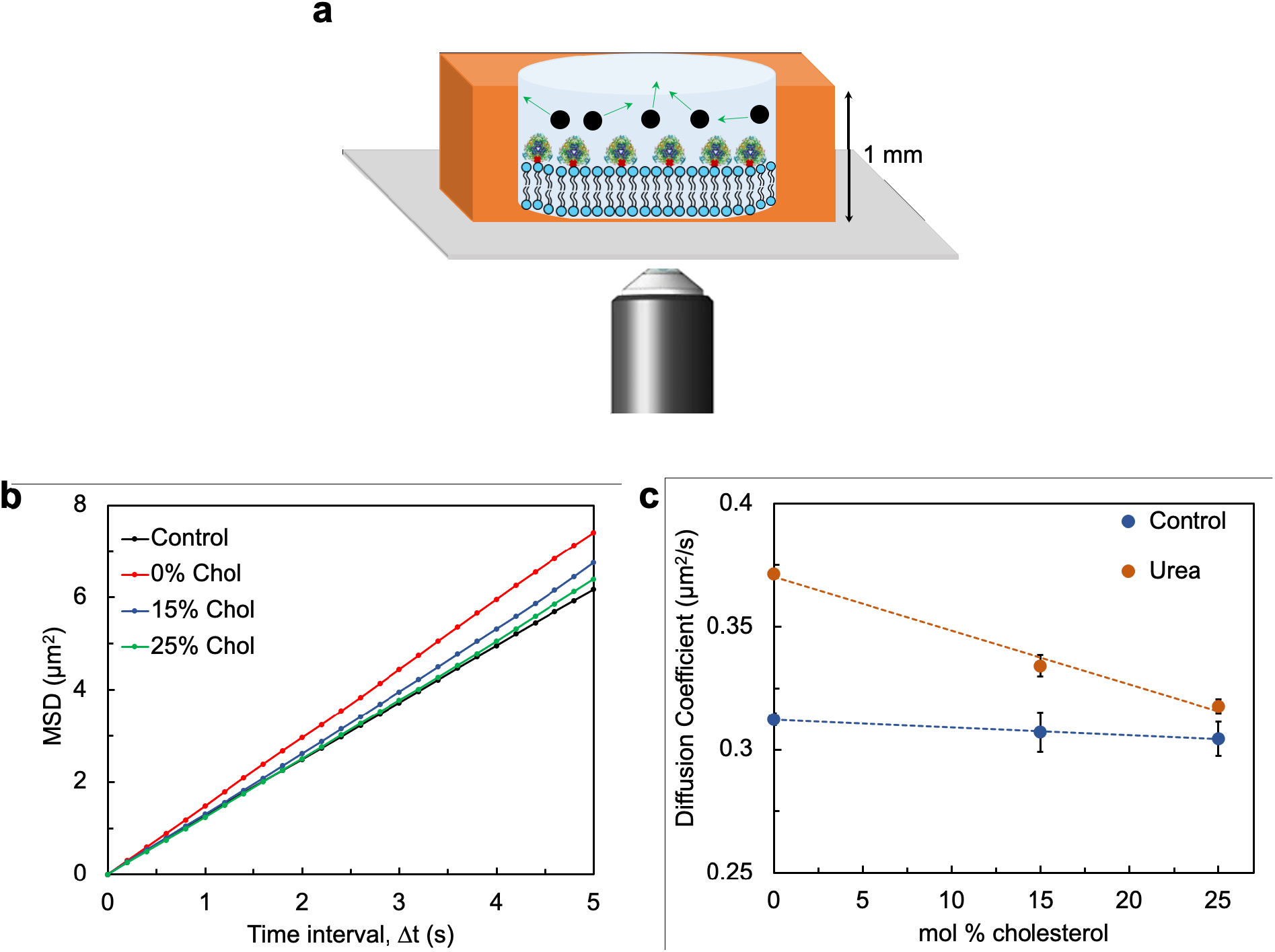
(a) Schematic representation of Supported Lipid Bilayer formed inside a hybridization chamber. 1 µm tracer particles are pipetted into the chamber to observe their diffusion behavior. The chamber is sealed at the top to prevent any air bubbles or unwanted flows (b) Mean Squared Displacement (MSD) of tracers upon catalysis of the bilayer at different cholesterol content. The control represents the average of tracer diffusion without catalysis at 0, 15, and 25 mol% cholesterol. The curves labelled 0, 15, and 25 mol% cholesterol represent tracer diffusion under these conditions in the presence of 200 mM urea. (c) Diffusion coefficient of tracer at 0, 15, and 25 mol% cholesterol with and without catalysis.

The passive tracers present on the bilayer undergo random diffusive motion. When the reactant, urea, was supplied to the bilayer, we observe an increase in the random diffusive motion of the tracers. This was quantified in terms of the diffusion coefficient of the particle as obtained from the MSD curves (**Figure 6b**). We observed that when the supported lipid bilayer had 0 mol% cholesterol there was an 18% enhancement in tracer diffusion. Increasing the cholesterol content in the bilayer progressively decreases the relative enhancement of tracer particles to 8% and 3% for 15 and 25% cholesterol respectively (**Figure 6c**). It is interesting to note here that while the relative enhancement in lipid diffusion remains constant across cholesterol concentrations, the enhancement in tracer diffusion decreases with increasing cholesterol content (**Figure 6c**). As shown in **Figure 3 and S2a**, the change in diffusion, ΔD (=D-D_o_) decreases with increasing cholesterol content even though the relative enhancement remains the same. We hypothesize that the smaller net increase in change in catalysis-induced lipid diffusivity with increasing cholesterol content is not sufficient to provide the force required for enhanced diffusion of the tracers. Previous theoretical studies have suggested that passive tracers on lipid bilayers can show higher diffusion due to membrane fluctuations.^31,32^ Our results provide the first experimental support for catalysis-induced enhancement of membrane fluctuations as an underlying mechanism.

### Directed transport of passive particles on asymmetric SLBs

As discussed, passive particles display enhanced diffusion on an SLB that is uniformly covered with enzymes. Now, we ask the question whether we can observe directional transport of particles by having an asymmetric distribution of enzymes on the SLB. We do this by using a PDMS mold to cover part of the glass substrate, allowing us to first pattern the remaining part of the coverslip with just POPC lipids. Removing the PDMS mold then enables us to attach POPC-Biotin-cap-DOPE (99:1) liposomes to just one area of the SLB. This allows the lipids with streptavidin-enzyme linkages to attach to only one region of the SLB, enabling us to form an enzyme-rich patch within the SLB.

In order to test directional transport of passive tracers around the enzyme-rich patch, we increased the size of the tracer particles to 2 µm and used a hybridization chamber height of 360 µm. This was done in order to reduce diffusive fluctuations and to prevent bulk convective flows, respectively. After the enzyme-rich patch was formed in the SLB (**Figure 7a**), BSA-coated 2 µm polystyrene tracer particles that were either added to buffer or to the reactant urea were pipetted into the hybridization chamber. The chamber was then sealed from the top using a glass coverslip to prevent air bubbles and unwanted disturbances that can cause convective flows in the sample. The movement of the tracer particles on the plane closest to the bilayer were then captured using an inverted optical microscope. We tracked the motion of the tracer particles in three different areas around the enzyme patch: right of the enzyme patch, bottom of the enzyme patch and top of the enzyme patch (**Figure S6**). A minimum of 15 particles (5 on each side of the enzyme patch) are tracked for control (buffer only) and for experiment with urea (in buffer). Their trajectories were used to calculate the average velocities of the tracer particles. We observe that on all three sides of the enzyme patch, the tracer particles move “outward” or away from the enzyme patch. As shown in **Figure 7b**, the average velocity of tracer particles is 0.3 µm/s outward from the patch. Note that while calculating the average velocity of tracer particles the positive direction is taken to be in the outward direction regardless of whether we are observing to the right, bottom, or top of the enzyme patch. **Figure 7c and d** show the Y positions of five tracer in the control and catalysis cases, respectively, when observed from above the enzyme patch. For each particle, the positions reported are relative to the initial position of the particle at time zero. The tracks of particles observed to the right and to the bottom of the patch are shown in **Figure S7**.

**Figure 7.**
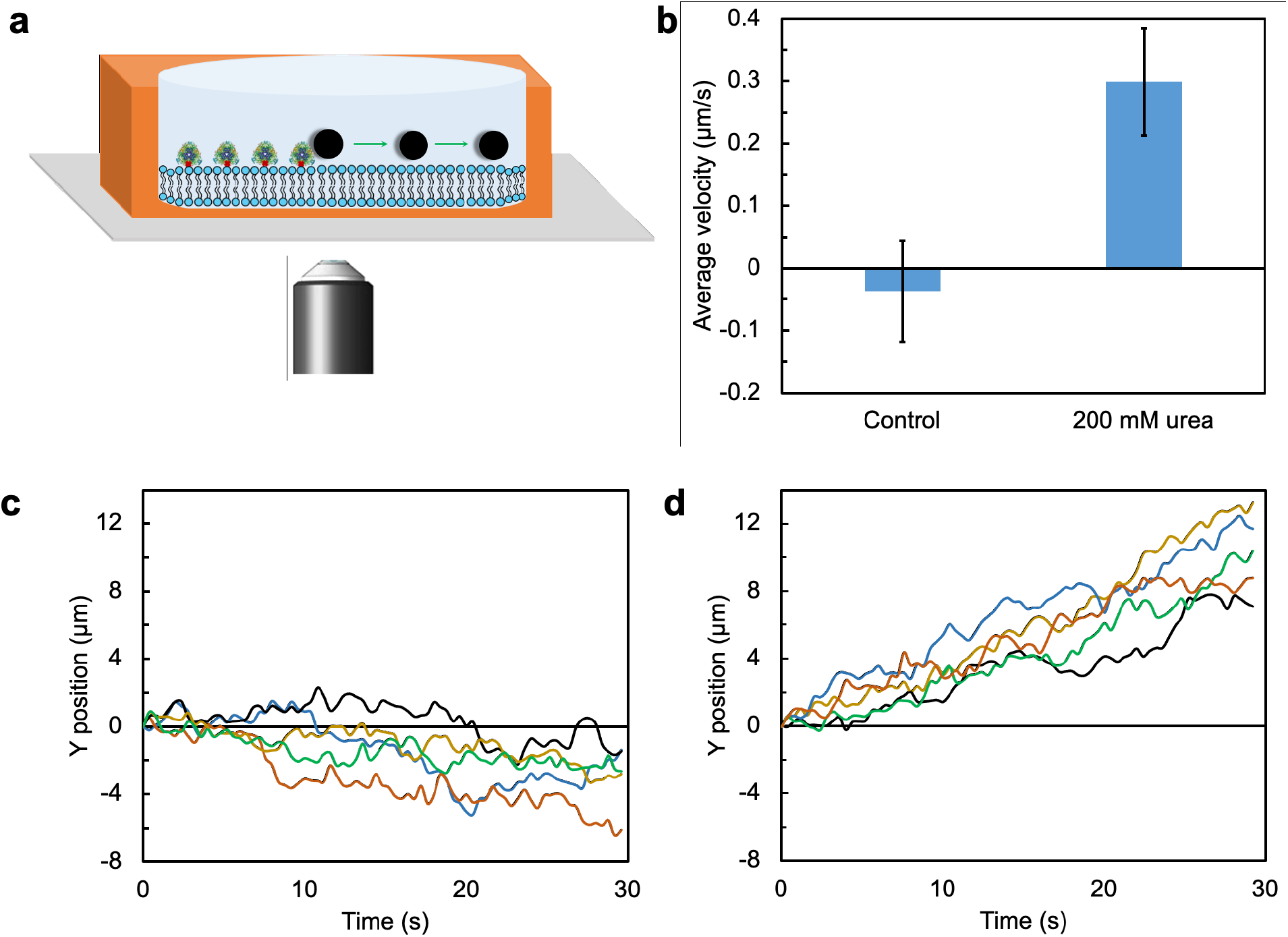
(a) Schematic of enzyme-rich patch formed inside a hybridization chamber. 2 µm particles were pipetted into the chamber to observe their directional transport behavior. The height of the hybridization chamber was 360 µm. (b) Average tracer particle velocities observed 100 µm away from the enzyme patch. The velocities were measured on three areas around the patch. Note that a positive velocity means that the particles are moving “outward” from the patch. (c) Tracks of five tracer particles recorded on one side of the enzyme patch in presence of 100 mM PBS buffer (d) Tracks of five tracer particles recorded on one side of the enzyme patch in presence of 200 mM urea.

A possible mechanism for the directional transport of tracers involves fluid pumping by catalytically-active enzymes. Previous experiments with enzymes-attached to solid substrates had shown that, during catalysis, immobilized enzymes create convective flows in the surrounding fluid.^33,34^ For urease, the direction of the flow is outward, away from the immobilized enzyme.^33^ Convective flow speeds are known to be a sensitive function of chamber height decreasing rapidly with decreasing chamber heigh.^33^ Indeed, when we decreased the height of the chamber from 360 to 120 µm, we observed that the directional transport of the tracers was quenched (**Figure S8**).

## Conclusion

In conclusion, we have demonstrated catalysis-induced enhanced diffusion of urease and hexokinase-attached lipids on SLB using fluorescence tagging and FRAP technique. We show that increasing the cholesterol content in the bilayer suppresses the overall diffusion but does not alter the relative enhancement in diffusion of the enzyme-attached lipids. The enhanced diffusive behavior of active lipids can be suppressed by attaching the enzyme to the SLB via a long and flexible linker. Using optical microscopy tracking, we have also demonstrated the first experimental evidence of catalysis-induced membrane fluctuations leading to the enhanced diffusion of passive tracers resting on the SLB. Additionally, by using active enzyme patches, we have demonstrated the directional transport of passive tracers on SLBs. This observation suggests a possible mechanism for the observed directional motility of influenza virus on cell surfaces.^35^ Our results constitute the first steps in understanding diffusion and transport in lipid membranes due to active, out-of-equilibrium processes that are the hallmark of living systems. More generally, our study demonstrates how active enzymes can be used to control diffusion and transport in confined 2-D environments and the design of motion-based sensing platforms.^36^

## Methods

### SLB formation

For the Fluorescence Recovery After Photobleaching (FRAP) experiments, liposomes of POPC and Biotin-cap-DOPE was prepared in a molar ratio of 99:1. Briefly, 100 µL of 500 µM liposome solution was pipetted out onto a glass slip with a circular PDMS well. The solution was allowed to fuse with the glass for 10 minutes. After which, it was washed multiple times with DI water to remove unattached excess liposomes. 50 µL of 1 µM Cy5 streptavidin was then added to the chamber for 10 minutes for its attachment onto the biotin heads of the SLB. The surface was then washed multiple times with DI water to remove any unattached streptavidin from the SLB. Biotinylated enzymes were then added to the PDMS well for 10 minutes to allow their attachment to the streptavidin attached SLB. After this step, the surface was washed with buffer multiple times to remove unattached excess enzymes from the surface. The surface of the SLB was scratched with a metal tweezer, to help with visualizing the surface of the lipid bilayer under confocal microscopy. The SLB was washed once again with buffer. The enzyme-attached SLBs were now ready for FRAP experiments. More details of the FRAP experiments are given in the SI.

### Optical microscopy analysis of passive tracers on SLB

Once the bilayer was prepared as mentioned above, we add 50 µl of the BSA-coated tracer particle suspension into the hybridization chamber. The chamber was then sealed with a glass coverslip to avoid air bubbles or unwanted disturbances that can cause convective flows. For the video scans, the objective was set at 50× and videos of tracers on the supported lipid bilayer were captured by optical microscope at 60 frames per second. We ensured that the videos were taken right at the surface of the bilayer by observing that there were some stuck particles in the frame. This told us that we were in the plane closest to the bilayer. The dimension of each frame corresponds to 152.9 µm along the x-axis and 114.07 µm along the y-axis which was set in the video tracker software while obtaining the trajectories of the tracer particles. A minimum of 10 particles were tracked and their trajectories were then used to calculate the MSD curves using MATLAB.

### Tracking directional transport of passive particles on SLB

In order to study the directional transport of passive tracers, SLBs with enzyme-rich patches were synthesized. We prepared two liposomes solutions that were used sequentially. One solution of liposomes was made of 100 mol% POPC lipids. The second liposome solution was with POPC and Biotin-cap-DOPE in a molar ratio of 99:1. We used a circular PDMS mold of 3 mm diameter and placed it inside the hybridization chamber. 100% POPC liposomes were first pipetted into the hybridization chamber so that the POPC SLB will form everywhere in the hybridization chamber except where the circular PDMS mold is placed. After allowing the liposomes to fuse onto the glass coverslip, the liposomes were washed to remove any unattached liposomes. The circular PDMS mold was then carefully removed and 50 µl of POPC-Biotin-cap-DOPE (99:1) was pipetted into the hybridization chamber. Following this, the chamber was thoroughly washed again to remove unattached liposomes. The procedure for the attachment of streptavidin and enzymes was same as mentioned above. The dimension of the hybridization chamber used for these experiments are 13 mm × 0.12 mm. We attached three of the spacers together to obtain a height of 0.36 mm (or 360 µm).

Once the bilayer was prepared, 100 µl of 200× diluted 2 µm sulphate-modified polystyrene microparticles was pipetted into the chamber either with reactant urea or with just the buffer. The chamber was then sealed with a glass coverslip to avoid air bubbles or unwanted disturbances that can cause convective flows. We then recorded the movement of tracers. The motion of the particles was recorded for 30 seconds at 60 frames per second. We recorded the motion of the tracers on three sides of the enzyme patch: right of the enzyme patch, bottom of the enzyme patch and top of the enzyme patch. A minimum of 15 particles were tracked and their trajectories were used to calculate the average velocities of the particles.

## Data Availability

The data that support the findings of this study are available from A. Sen upon reasonable request.

## Acknowledgments

We gratefully acknowledge funding of the research by the Air Force Office of Scientific Research FA9550-20-1-0393. We thank Dr. Liqiang Ren whose MATLAB code was used to calculate MSD values of tracer particles.

## Author contribution

The work was conceived by A. Sen and A. Somasundar. A. Somasundar performed the experiments. N. Mandal did the calculations of two-dimensional viscosity and friction coefficient. All authors contributed to the discussion of results and writing of the manuscript.

